# Dietary vitamin B12 regulates chemosensory receptor gene expression via the MEF2 transcription factor in *Caenorhabditis elegans*

**DOI:** 10.1101/2021.10.02.462191

**Authors:** Aja McDonagh, Jeannette Crew, Alexander M. van der Linden

**Affiliations:** Department of Biology, University of Nevada, Reno, Nevada, 89557

**Keywords:** diet, vitamin B12, *Caenorhabditis elegans*, *srh-234*, chemosensory receptor gene, ADL, sensory neurons, plasticity

## Abstract

Dynamic changes in chemoreceptor gene expression levels in sensory neurons is one strategy that an animal can use to modify their responses to dietary changes. However, the mechanisms underlying diet-dependent modulation of chemosensory gene expression are unclear. Here, we show that the expression of the *srh-234* chemoreceptor gene localized in a single ADL sensory neuron type of *C. elegans* is downregulated when animals are fed a *Comamonas* bacterial diet, but not on an *E. coli* diet. Remarkably, this diet-modulated effect on *srh-234* gene expression levels is dependent on the micronutrient vitamin B12 endogenously produced by *Comamonas* bacteria. Excess propionate and genetic perturbations in the canonical and shunt propionate breakdown pathways are able to override the repressing effects of vitamin B12 on *srh-234* expression. The vitamin B12-mediated regulation of *srh-234* expression levels in ADL requires the MEF-2 transcription factor, providing a potential mechanism by which dietary vitamin B12 may transcriptionally tune individual chemoreceptor genes in a single sensory neuron type, which in turn may change animal responses to biologically relevant chemicals in their diet.

## INTRODUCTION

Animals receive dietary inputs from their environment and their internal metabolic state, which allows them to modify their chemosensory response properties and behavioral outcomes (Sengupta 2012). One strategy that animals can use to trigger long term changes in behavioral outcomes is by dynamically changing the expression of individual chemoreceptor genes present in chemosensory neurons. These dynamic changes in chemoreceptor gene expression in response to food and internal feeding state is observed in different invertebrate systems and play pivotal roles in their ability to seek food and reproduce (Fox et al. 2001; Hallem et al. 2004; Rinker et al. 2013; Ryan et al. 2014; Khan et al. 2021; Taparia et al. 2017), but the mechanisms controlling this plasticity in chemoreceptor gene expression are unclear.

The nematode *C. elegans* is an excellent model organism to study interactions between an animal and its dietary sources (Zhang et al. 2017; Yilmaz and Walhout 2014). *C. elegans* is a bacterivore, making it easy to expose *C. elegans* to different bacterial strains to study their effects on organismal health and physiology. Bacterially-derived factors affect *C. elegans* in various ways; for instance, pathogenic factors are sensed by chemosensory neurons and trigger avoidance behaviors (Pradel et al. 2007; Meisel et al. 2014), while other bacterially-derived factors are innocuous and contribute to physiology and development (Coolon et al. 2009; Gracida and Eckmann 2013). Recent work demonstrated that vitamin B12 obtained by *C. elegans* through its bacterial diet is an important nutritional factor in developmental growth and physiology of *C. elegans* (MacNeil et al. 2013). The vitamin B12 status of *C. elegans* can be easily assessed with help of the *acdh-1p::gfp* reporter, which is expressed in response to propionate accumulation resulting from B12 deficiency (Watson et al. 2013; Watson et al. 2014; Watson et al. 2016). When fed a vitamin B12-deficient *E. coli* OP50 diet, *acdh-1* is highly expressed in animals, whereas *acdh-1* is lowly expressed when grown on the vitamin B12-producing *Comamonas* DA1877 diet. The effects of these bacterial diets on *acdh-1* promoter activity have led to important insights into the vitamin B12-dependent and independent propionate breakdown pathways.

*C. elegans* is also an ideal organism to study the plasticity in expression levels of individual chemosensory receptor genes in response to external and internal signals (Vidal et al. 2018; Gruner and van der Linden 2015). Our prior study showed that the expression levels of the *srh-234* chemoreceptor gene in the ADL sensory neuron type is regulated by starvation. This starvation-mediated modulation of *srh-234* expression levels is dependent on sensory inputs into ADL neurons perceiving food presence, and circuit inputs from RMG interneurons that are electrically connected to ADL perceiving internal state of starvation signals (Gruner et al. 2014). Circuit inputs from RMG into ADL regulating *srh-234* required the NPR-1 neuropeptide receptor acting in RMG, as well as insulin signals from other tissues acting on the DAF-2 insulin receptor in ADL (Gruner et al. 2014). In addition, starvation-mediated regulation of *srh-234* expression levels in ADL is regulated by both cell- and non-cell-autonomous transcriptional mechanisms involving basic helix-loop-helix (bHLH) factors, including HLH-30 and MXL-3 acting in the intestine, and HLH-2/3 acting together with the MEF-2 factor in ADL neurons (Gruner et al. 2016). Together, these findings demonstrated that expression of the *srh-234* chemoreceptor gene in a single ADL sensory neuron type of *C. elegans* is regulated by multiple transcriptional modules, and revealed a neuron-to-intestine connection involving insulin signals in the modulation of chemoreceptor genes as a function of the *C. elegans* feeding state (Gruner and van der Linden 2015).

In this study, we discovered that feeding *C. elegans* vitamin B12-producing *Comamonas* bacteria regulates the expression levels of the *srh-234* chemoreceptor gene in ADL neurons. We show that *srh-234* gene expression is strongly downregulated in ADL when animals are fed a high vitamin B12 diet of *Comamonas* DA1877 bacteria relative to a low vitamin B12 diet of *E. coli* OP50 bacteria. This dietary effect of vitamin B12 on *srh-234* in ADL appears to be distinct from the starvation response we previously reported (Gruner et al. 2014). Mutant bacteria of *Comamonas* deficient in vitamin B12 production indicated that *Comamonas*-supplied vitamin B12 regulates *srh-234* expression levels in ADL. The repressing effects of vitamin B12 on *srh-234* can be suppressed by propionate supplementation and genetic perturbations in the canonical and shunt propionate breakdown pathways. The effects of vitamin B12 on *srh-234* expression is likely through food ingestion rather than directly sensing B12. Vitamin B12-mediated downregulation of *srh-234* is dependent on the MEF-2 transcription factor. Together, these findings reveal that bacterially-derived vitamin B12 turn individual chemoreceptor genes on and off at the level of transcription in sensory neurons that may inform our understanding of how animals fine-tune their chemosensory responses to biologically relevant chemicals in their diet.

## MATERIAL AND METHODS

### *C. elegans* strains and growth conditions

Strains used in this study were: wild-type N2 *C. elegans* variety Bristol, RB1774 *pcca-1(ok2282)*, VC1307 *pccb-1(ok1686)*, VC1011 *acdh-1(ok1489)*, RB2572 *hphd-1(ok3580)*, RB755 *metr-1(ok521)*, NYL2498 *mrp-5(yad138)*, JIN1375 *hlh-30(tm1978)*, and KM134 *mef-2(gv1)*. Transgenic strains used in this study were: VDL3 *oyIs56*[*srh-234p::gfp, unc-122p::rfp*], VDL497 *sanEx497*[*sre-1p::gfp, rol-6*], VDL494 *sanEx494*[*sre-1p(+MEF2)::gfp, rol-6*], and VL749 *wwIs24*[*acdh-1p::gfp, unc-119(+)*]. Animals were cultivated at 20°C on the surface of Nematode Growth Media (NGM) agar. Unless specified otherwise, animals were fed *E. coli* OP50 as the primary food source (Brenner 1974). Genotypes used in this study were confirmed by PCR (for example, identifying deletions), or by sequencing a PCR product (for example, identifying single nucleotide changes).

### Bacterial strains and growth conditions

Bacterial strains used in this study were: *E. coli* OP50, *E. coli* HT115 (DE3), *E. coli* HB101, *E. coli* BW25113, *E. coli* Δ*tonB* JW5195, *Comamonas aq*. DA1877, *Comamonas aq*. Δ*cbiA* and Δ*cbiB* mutants. Bacterial cultures were grown under standard conditions in Luria Broth (LB) media until the Optical Density (OD) 600 reached approximately 0.6. *Comamonas* mutants were cultured in the presence of 100 μg/ml streptomycin plus 20 μg/ml gentamycin as a selection marker. Presence of these antibiotics did not alter the levels of *srh-234p::gfp* expression.

### Measurement and quantification of *gfp*-reporter expression levels

Animals carrying chemoreceptor::*gfp* reporter genes (i.e., *srh-234, sre-1*) were cultivated at 20°C on NGM plates seeded with *E. coli* OP50 as the bacterial food source unless indicated otherwise. Gravid adults were transferred to assay plates and removed after laying eggs. The eggs were then allowed to develop to adults. The increased rate of development when fed *Comamonas* DA1877 was accounted for, and levels of promoter::*gfp* expression of adult animals were then imaged and measured under a microscope equipped with epifluorescence as previously described (Gruner et al. 2014; Gruner et al. 2016). Briefly, we mounted animals on 2% agarose pads containing 10 mM levamisole, and visualized them on a Leica DM5500 compound microscope equipped with epifluorescence and a Hamamatsu CCD-camera. Microscope and camera settings were kept constant between images of different genotypes and conditions used, unless indicated otherwise. The mean pixel intensity of *gfp* fluorescence in the entire cell-body of ADL was quantified using Volocity software (version 6.3). Prior to measurement, images of ADL cell-bodies were cropped for promoter-*gfp* expression level analysis.

### Analysis of *srh-234p::gfp* expression

To analyze *srh-234* expression in mixed bacterial diets, animals carrying the *srh-234p::gfp* reporter were exposed to mixed set ratios, i.e., 1:1, 9:1, and 99:1 ratio of *E. coli* OP50 to *Comamonas aq*. DA1877. To prepare plates, liquid bacterial cultures of OP50 and DA1877 were grown overnight at 37°C in LB broth, and diluted or concentrated to the same OD600. Bacteria were seeded onto peptone-free NGM agar plates to minimize bacterial growth. Adults expressing the *srh-234p::gfp* reporter were transferred to plates and removed after eggs were laid. Eggs were allowed to develop to adulthood in the presence of the mixed bacterial diets, and *srh-234p::gfp* expression levels were measured and quantified as described above.

To analyze *srh-234* expression in the presence of exogenous vitamin B12 and propionic acid (aka propionate), animals carrying the *srh-234p::gfp* reporter were transferred to NGM plates seeded with *E. coli* OP50 supplemented with or without vitamin B12 (methylcobalamin or MeCbl, Sigma, Cat #13422-55-4; adenosylcobalamin or AdoCbl, Sigma, Cat #13870-90-1) and propionic acid (Sigma, Cat #79-09-4). Stocks were made in either ethanol (for MeCbl) and water (for AdoCbl and Propionic acid) to the maximum soluble concentration. Vitamin B12 and propionic acid was diluted to a final 64 nM and 40 mM concentration, respectively, in NGM agar prior to plate pouring. For *E. coli* OP50 supplementation assays with increasing MeCbl concentrations, we created a dilution series from a 1 mM MeCbl stock. To confirm vitamin B12 action, *acdh-1p::gfp* reporter animals were used as a control in parallel to the *srh-234p::gfp* expression analysis.

For bacterial olfactory assays, *srh-234p::gfp* reporter animals were exposed to either *E. coli* OP50 or *Comamonas* DA1877 bacteria seeded on a NGM agar square placed on the inside of a petri dish lid. For the quadrant petri dish assay, NGM plates were seeded in each quadrant with either OP50 or DA1877 diets (**Fig. S4**). *srh-234p::gfp* reporter animals were then transferred to a single quadrant of the plate allowing only a single diet for food ingestion, while allowing olfactory cues of the surrounding diets.

For generating inedible food, *Comamonas* DA1877 bacteria were treated with the antibiotic aztreonam (Sigma, Cat #78110-38-0). Briefly, DA1877 bacteria were grown in LB to log phase at 37°C with shaking. Cultures were mixed with aztreonam to a final concentration of 10 μg/ml for an additional three hours with minimal shaking to prevent bacterial shearing. Aztreonam-treated bacteria were spread onto the NGM agar plates and immediately dried and used the same day, because the septum inhibitory effects of aztreonam are short lived. *srh-234p::gfp* reporter animals were then transferred as young adults to plates containing aztreonam-treated DA1877.

### Dye-filling of ADL sensory neurons

A stock dye solution containing 5 mg/μl red fluorescent lipophilic dye DiI (Sigma, Cat #41085-99-8) was diluted in M9 buffer by 10,000 times for optimal signal intensity. Animals carrying the *srh-234p::gfp* reporter were soaked in DiI for one hour and then rinsed with M9 buffer twice. Stained animals were recovered for one hour on NGM plates seeded with either *E. coli* OP50 or *Comamonas* DA1877 before examination of dye-filled ADL neurons with a Leica DM5500 microscope equipped with epifluorescence.

### Statistical analysis

All results are expressed as means with 95% confidence intervals. Data sets were first analyzed for Gaussian distribution using a normality test (alpha =0.05, p>0.05) using either the Shapiro-Wilk test or D’Agostino and Pearson normality test to determine whether a parametric or non-parametric statistical test should be performed. Statistical comparisons made for two groups include an unpaired *t*-test (parametric) or the Mann-Whitney *t*-test (non-parametric). For more than two groups, the ordinary one-way ANOVA (parametric) or the Kruskal-Wallis test (non-parametric) was used followed by a posthoc multiple comparisons test. Specific statistical tests and *p*-values are reported in the Figure legends. All data were graphed and analyzed using Graphpad Prism 9 software.

## RESULTS

### Expression of *srh-234* is downregulated when animals are fed a *Comamonas* diet

To study how bacterial diet regulates chemoreceptor gene expression levels in *C. elegans*, we used the candidate *srh-234* chemoreceptor gene specifically expressed in a single sensory neuron type, ADL. We previously found that *gfp* expression driven by only 165 bp *cis*-regulatory sequence of *srh-234* (referred to as *srh-234p::gfp*) is rapidly (<1 hr) downregulated in starved animals (Gruner et al. 2014). While testing the *srh-234p::gfp* reporter in different bacterial diets, we observed that animals fed a *Comamonas* DA1877 diet downregulate *srh-234* expression in ADL neurons similar in response to starvation; that is *srh-234p::gfp* expression levels in adult animals is strongly reduced when fed a *Comamonas* DA1877 diet compared to a *E. coli* OP50 diet (**Fig. 1A**). This *Comamonas*-mediated downregulation of *srh-234* expression levels is rapid as adults raised on *E. coli* OP50 and then transferred to a DA1877 diet reduce *srh-234p::gfp* expression in ADL neurons by 50% after 2 hours (**Fig. S1A**). Animals fed other *E. coli* diets such as the K12/B-type hybrid HB101 strain, and the K12-type HT115 strain commonly used in *C. elegans* research showed a *srh-234* expression phenotype intermediate to that of *E. coli* OP50 and *Comamonas* DA1877 diets (**Fig. S1B**).

**Fig. 1:**
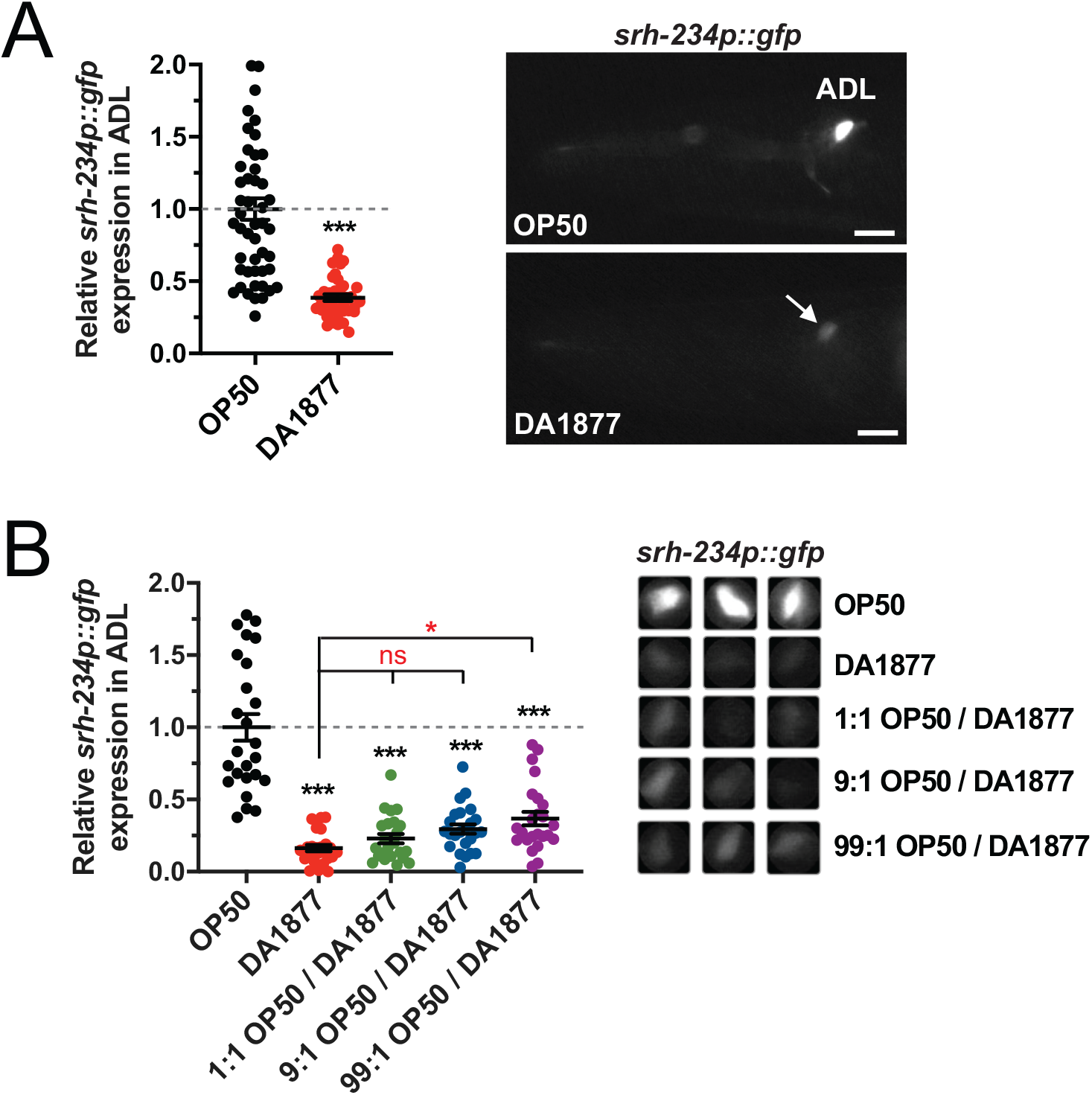
Expression of *srh-234* is downregulated by a *Comamonas* DA1877 diet. **(A)** Relative expression levels of *srh-234p::gfp* in the ADL cell body of adults fed either an *E. coli* OP50 diet or a *Comamonas* DA1877 diet. Adult animals containing stably integrated copies of a *srh-234p::gfp* transgene (*oyIs56*) were examined at the same exposure time on both diets. Images are lateral views of the ADL sensory neuron. Scale is 15 μm. Data are represented as the mean ± SEM (n>38 animals for each diet). *** *p*<0.001 by an unpaired 2-tailed *t*-test. **(B)** Relative expression levels of *srh-234* in the ADL cell body of adult animals fed each of the indicated diets. OP50: *E. coli*; DA1877 *Comamonas*; 1:1, 9:1 and 99:1 refers to the dilution of *Comamonas* DA1877 in *E. coli* OP50. Bacteria were seeded on peptone-free plates to prevent bacterial growth (see Material and Methods). Right panel: Representative cropped images of *srh-234p::gfp* expression in the ADL cell body with the indicated diets and dilutions. Images were acquired at the same exposure time. Data are represented as the mean ± SEM (n>24 animals for each condition). *** indicates values that are different from wild-type animals fed on *E. coli* OP50 at *p*<0.001 by a Kruskal-Wallis with Dunn multiple-comparisons test.

The dietary effect of *Comamonas* DA1877 on *srh-234* expression appears to be distinct from the starvation response, because mixing the *E. coli* OP50 diet with *Comamonas* DA1877 diet 1:1 resulted in animals in which *srh-234* expression levels remained strongly reduced similar to starvation (**Fig. 1B**). Moreover, smaller concentrations of *Comamonas* DA1877 by diluting it in *E. coli* OP50 (i.e., 9:1 and 99:1 OP50/DA1877) was sufficient to strongly reduce *srh-234* expression. Others have reported that *Comamonas* DA1877 bacteria are not a nutrient-poor diet for *C. elegans* (Shtonda and Avery 2006; MacNeil et al. 2013), suggesting that *Comamonas* may generate a bacterial signal that regulates *srh-234* expression levels. This dietary effect of *Comamonas* on *srh-234* may be specific since expression of another ADL-specific *sre-1* chemoreceptor is not affected (**Fig. S1C**). Since we previously showed that altered sensory (i.e. cilia, dendrites) inputs into ADL neurons can dramatically reduce *srh-234p::gfp* expression levels (Gruner et al. 2014), it remains possible that *Comamonas* affects the integrity of ADL neurons; however, animals show normal dye-filling (100% of animals dye-fill, n>20) and a normal ADL morphology determined by *sre-1p::gfp* expression when fed with the *Comamonas* DA1877 diet (**Fig. S1D**). Together, these results suggest that in addition to starvation, a dilutable bacterial metabolite produced by *Comamonas* bacteria regulates *srh-234* expression levels in ADL neurons.

### Vitamin B12 produced by *Comamonas aq*. represses *srh-234* expression

The strain *Comamonas* DA1877 produces the dilutable metabolite vitamin B12, while the *E. coli* OP50 strain is not able to synthesize vitamin B12 (Watson et al. 2014). To test the hypothesis that vitamin B12 downregulates *srh-234* expression levels in ADL neurons, we examined *C. elegans* animals fed a *E. coli* OP50 diet supplemented with two biologically active and interconvertible forms of vitamin B12, adenosylcobalamin (AdoCbl) and methylcobalamin (MeCbl). We found that animals fed an *E. coli* OP50 diet supplemented with either 64 nM AdoCbl or MeCbl was sufficient to strongly reduce *srh-234p::gfp* expression in ADL neurons (**Fig. 2A**), suggesting that vitamin B12 represses the expression of *srh-234*. Moreover, supplementing *E. coli* OP50 with increasing concentrations (nM doses) of MeCbl resulted in a dose-dependent reduction of *srh-234p::gfp* expression (**Fig. S2A**), which fits with our observation that diluting *Comamonas* into the *E. coli* diet is sufficient to reduce *srh-234* expression levels (**Fig. 1B**). As a control, we found similar dose-dependent effects of MeCbl using the *acdh-1p::gfp* reporter (**Fig. S2B**), which is known to be downregulated in the intestine when fed the vitamin B12-producing *Comamonas* bacteria or when fed *E. coli* OP50 supplemented with vitamin B12 (Watson et al. 2014; MacNeil et al. 2013). These results are also consistent with the observed *srh-234* expression phenotype of animals raised on *E. coli* HT115 and HB101 diets (**Fig. S1B**), which have higher vitamin B12 levels compared to the *E. coli* OP50 diet (Revtovich et al. 2019). Thus, vitamin B12 supplementation to an *E. coli* diet can repress the expression of *srh-234* in ADL.

**Fig. 2:**
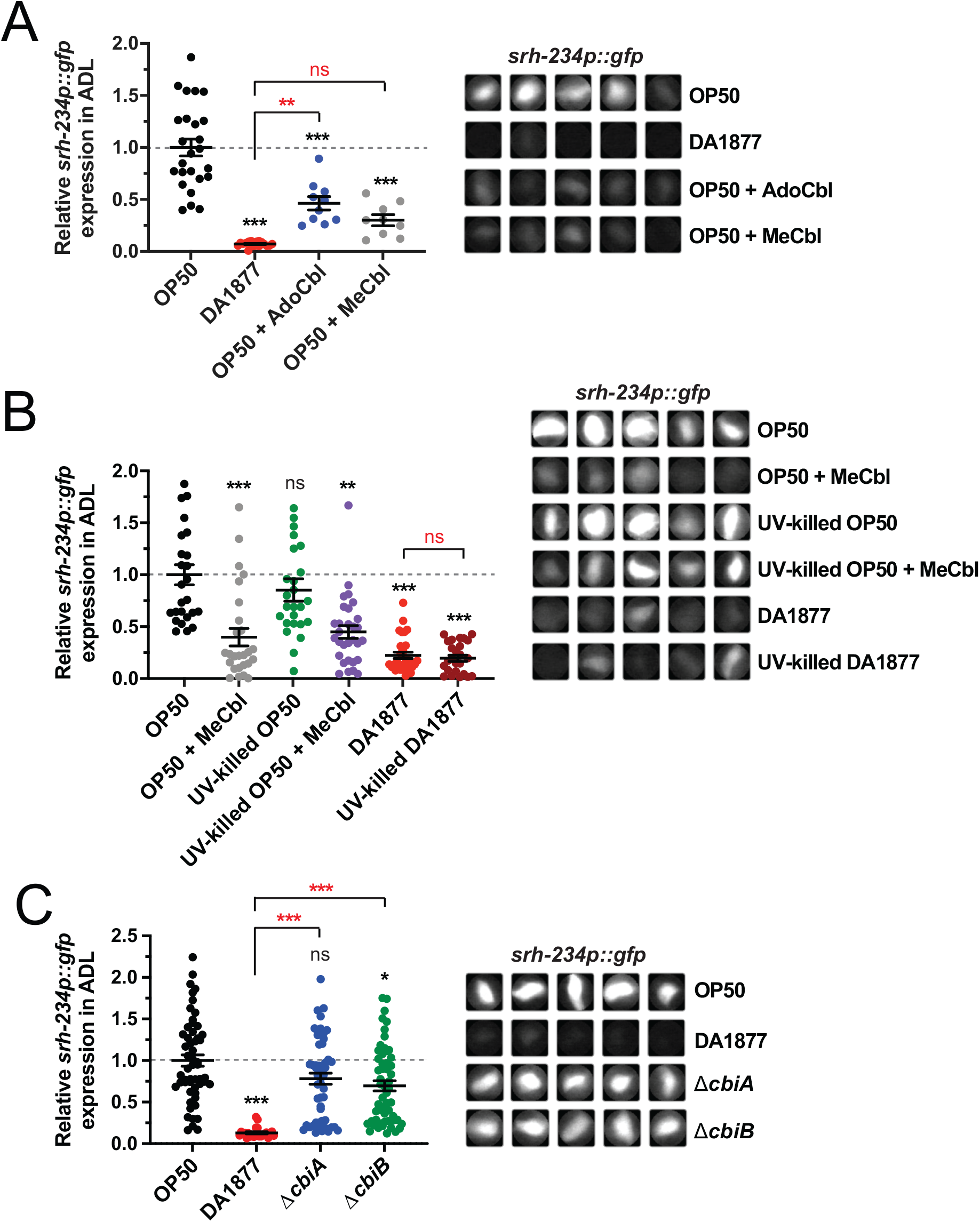
Vitamin B12 produced by *Comamonas* is required to downregulate *srh-234* expression. **(A)** Relative expression of *srh-234p::gfp* in the ADL cell body of OP50-fed adult animals supplemented with either AdoCbl or MeCbl compounds at a 64 nM final concentration. Data are represented as the mean ± SEM (n>9 animals for each condition). ***, ** indicates values that are different from wild-type fed on *E. coli* OP50 or *Comamonas* DA1877 at *p*<0.001 and *p*<0.01, respectively, by a one-way ANOVA with Tukey multiple-comparisons test. **(B)** Relative *srh-234* expression in the ADL cell body of animals fed either live or UV-irradiated killed *E. coli* OP50 and *Comamonas* DA1877 diets. Data are represented as the mean ± SEM (n>26 animals for each condition). ***, ** indicates values that are different from wild-type animals fed on *E. coli* OP50 or *Comamonas* DA1877 at *p*<0.001 and *p*<0.01, respectively by a Kruskal-Wallis with Dunn multiple-comparisons test. **(C)** Relative expression of *srh-234* in the ADL cell body of adults fed the *Comamonas* mutant strains Δ*cbiA* and Δ*cbiB* defective in producing vitamin B12 diets compared to *E. coli* OP50 and *Comamonas* DA1877. Data are represented as the mean ± SEM (n=24-56 animals for each diet). *** *p*<0.001, ** *p*<0.01 by a Kruskal-Wallis with Dunn multiple-comparisons test. **(A-C)** Right panels: Representative cropped images of *srh-234p::gfp* expression in the ADL cell body with the indicated compounds, genotypes and/or conditions. Images were acquired at the same exposure time. ns, not significant.

The vitamin B12-mediated reduction in *srh-234* expression levels in ADL could be explained by the fact that additional vitamin B12 added to *E. coli* may alter the metabolism of these bacteria by, for instance, decreasing the production of a toxic bacterial metabolite. Alternatively, *E. coli* may modify or metabolize vitamin B12 by creating a secondary by-product which in turn could reduce *srh-234* expression levels. To distinguish between these possibilities, we fed animals expressing the *srh-234p::gfp* reporter either live or ultraviolet (UVC)-killed *E. coli* OP50 bacteria in the presence of vitamin B12 and compared their *srh-234* expression levels. While UVC-killed bacteria of *E. coli* OP50 in the absence of vitamin B12 did not significantly alter *srh-234* expression (**Fig. 2B**), we found that *srh-234p::gfp* expression in the presence of vitamin B12 (64 nM MeCbl) is repressed equally well when supplemented to either live or UVC-killed *E. coli* OP50 bacteria (**Fig. 2B**). Similarly, UVC-killed *Comamonas* DA1877 did not affect the *srh-234p::gfp* expression levels. These findings suggest that the effects of vitamin B12 on *srh-234* gene expression levels do not appear to depend on *E. coli* modification or its metabolism.

To further test whether *Comamonas*-supplied vitamin B12 regulates *srh-234* expression in ADL neurons, we took advantage of mutant strains of *Comamonas* bacteria that are deficient in vitamin B12 production, and also fail to reduce expression levels of the *acdh-1p::gfp* intestinal reporter (**Fig. S2C**). We found that transposon mutations in genes of the vitamin B12 biosynthetic pathway of *Comamonas* DA1877 Δ*cbiA* and Δ*cbiB* that produce little or no vitamin B12 in these bacteria (Watson et al. 2014), fail at least in part to reduce *srh-234p::gfp* expression in ADL as observed in DA1877-fed animals (**Fig. 2C**). Together, these results suggest that vitamin B12 synthesized by *Comamonas* bacteria regulates the expression of *srh-234* in ADL neurons.

### Propionate overrides the repressing effects of vitamin B12 on *srh-234* expression

Since the balance between vitamin B12 and propionyl-CoA levels involved in propionate breakdown (**Fig. 3A**) has been reported to control promoter activity of the *acdh-1* gene (Watson et al. 2016), we next tested whether propionate can also regulate *srh-234* expression levels. We found that animals fed on an *E. coli* OP50 diet in the presence of vitamin B12 restored *srh-234p::gfp* expression in ADL neurons to near wild-type levels when supplemented with excess propionate (**Fig. 3B**). Feeding animals an *E. coli* OP50 diet supplemented with propionate alone did not significantly alter *srh-234p::gfp* expression levels in ADL (**Fig. 3B**). Thus, similar to the *acdh-1* promoter, excess propionate can override the repressing effects of vitamin B12 on *srh-234* expression in ADL neurons.

**Fig. 3:**
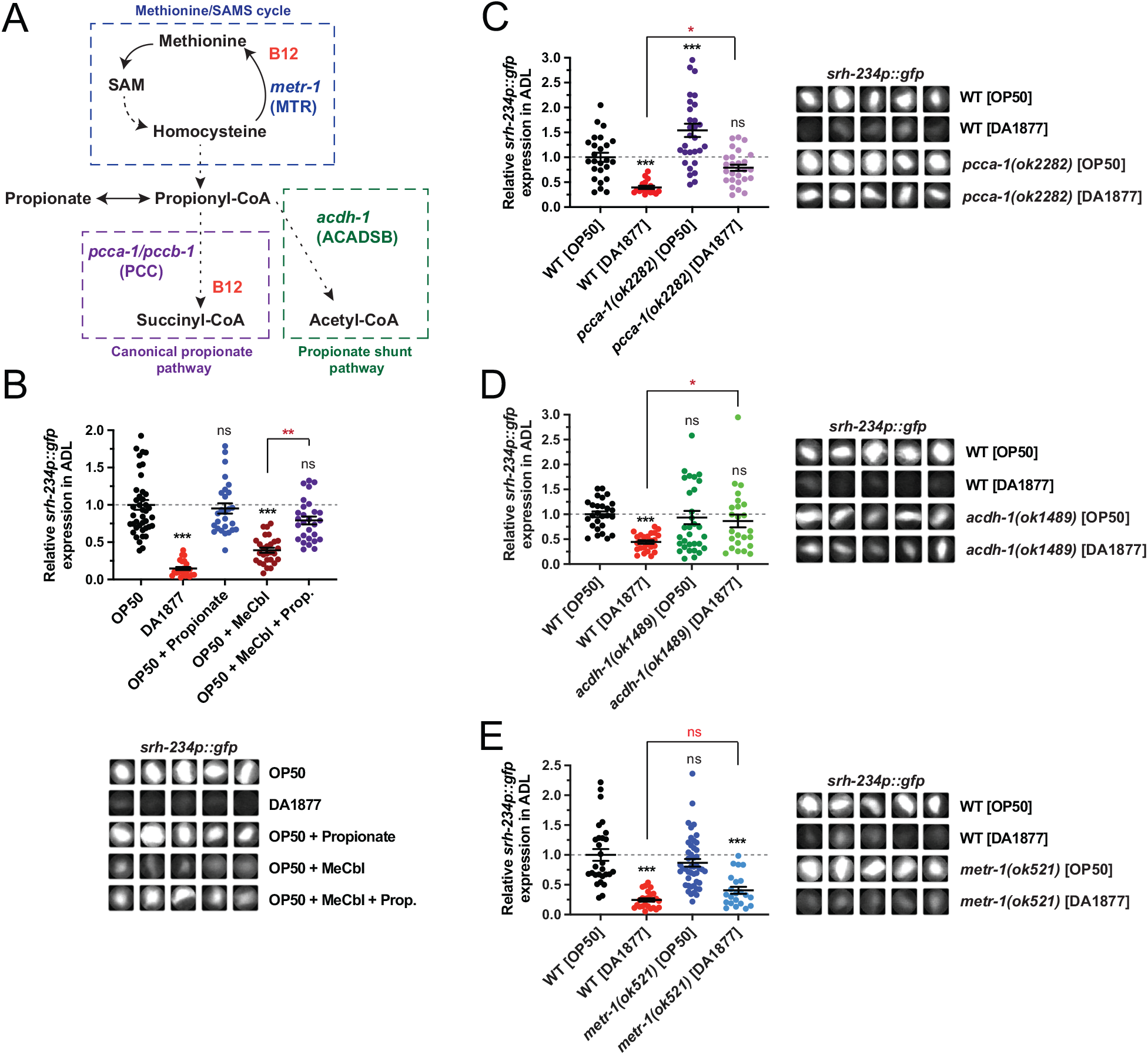
Downregulation of *srh-234* expression by vitamin B12 is reversed by propionate accumulation. **(A)** Schematic of the *C. elegans* methionine/SAMs cycle (blue dotted box), canonical propionate breakdown pathway (purple dotted box), and the propionate shunt metabolic pathway (green dotted box). **(B)** Relative expression of *srh-234p::gfp* in the ADL cell body of adults fed an *E. coli* OP50 diet supplemented with excess propionate (40 mM final concentration) and/or MeCbl (64 nM final concentration). Data are represented as the mean ± SEM (n=24-42 animals for each condition). ***, ** indicates values that are different from wild-type fed on *E. coli* OP50 or *Comamonas* DA1877 at *p*<0.001 and *p*<0.01, respectively, by a one-way ANOVA with Tukey multiple-comparisons test. **(C-E)** Relative expression of *srh-234p::gfp* in the ADL cell body of adult animals with mutations in the canonical propionate breakdown pathway **(C)**, the propionate shunt breakdown pathway **(D)**, and the methionine/SAM cycle **(E)** fed either *E. coli* OP50 or *Comamonas* DA1877 diets. Data are represented as the mean ± SEM. n>18 animals for each diet and genotype. **(C)** ***, * indicates values that are different from wild-type fed on *E. coli* OP50 or *Comamonas* DA1877 at *p*<0.001 and *p*<0.05, respectively, by a one-way ANOVA with Tukey multiple-comparisons test. **(D-E)** *** *p*<0.001, * *p*<0.05 by a Kruskal-Wallis with Dunn multiple-comparisons test. **(B-E)** Images were acquired at the same exposure time. ns, not significant.

Low vitamin B12 diets such as *E. coli* OP50 or genetic perturbation of the canonical propionate breakdown pathway leads to propionate accumulation and the transcriptional activation of the propionate shunt pathway (Watson et al. 2013; Watson et al. 2014; Watson et al. 2016). Since vitamin B12 fails to fully reduce *srh-234* expression levels in ADL neurons in the presence of excess propionate, we next tested whether propionate buildup due to genetic perturbations in the canonical and shunt propionate breakdown pathways (**Fig. 3A**) also lead to changes in *srh-234* promoter activity. As expected, we found that animals reduce *srh-234p::gfp* expression when fed the vitamin B12-producing *Comamonas* DA1877, but not in those animals that carry mutations in the first step of the canonical propionate pathway, *pcca-1* and *pccb-1* (**Fig. 3C, S3A**) or in propionate shunt pathway genes, *acdh-1* and *hphd-1* (**Fig. 3D, S3B**). Interestingly, *srh-234p::gfp* expression is slightly increased in both *pccb-1* and *pcca-1* mutants fed on the low vitamin B12 *E. coli* OP50 diet compared to wild-type, possibly in response to a further accumulation of propionate. Mutations in the methionine/SAM cycle gene, *metr-1*, did not show significant effects on *srh-234p::gfp* expression in ADL compared to wild-type when fed on *Comamonas* DA1877 (**Fig. 3E**). Together, these results suggest that *srh-234* expression levels in ADL neurons are repressed by dietary-supplied vitamin B12 and activated by propionate levels.

### Dietary-supplied vitamin B12 reduces *srh-234* expression through food ingestion

We previously showed that *srh-234* expression is dependent on starvation associated with a decreased food ingestion of *E. coli* OP50, as well as sensory inputs into ADL neurons associated with a decreased presence of OP50 food (Gruner et al. 2014). To test whether vitamin B12 can act as a volatile olfactory chemical to alter levels of *srh-234* expression in ADL, we decided to expose animals expressing *srh-234p::gfp* to NGM agar plates seeded with *E. coli* OP50 bacteria that were covered with petri-dish lids containing NGM agar squares soaked with a 1 mM concentration of vitamin B12 (MeCbl) placed above the animals (**Fig. 4A**). In addition, we exposed worms to *Comamonas* DA1877 which they cannot eat or touch, while feeding *E. coli* OP50, and vice versa (**Fig. S4A-B**). In both assays, we found that expression levels of *srh-234* in ADL neurons was not significantly altered when exposed to *Comamonas* bacteria or vitamin B12, suggesting that vitamin B12 likely does not act as an olfactory chemical cue released by bacteria to regulate *srh-234* expression in ADL.

**Fig. 4:**
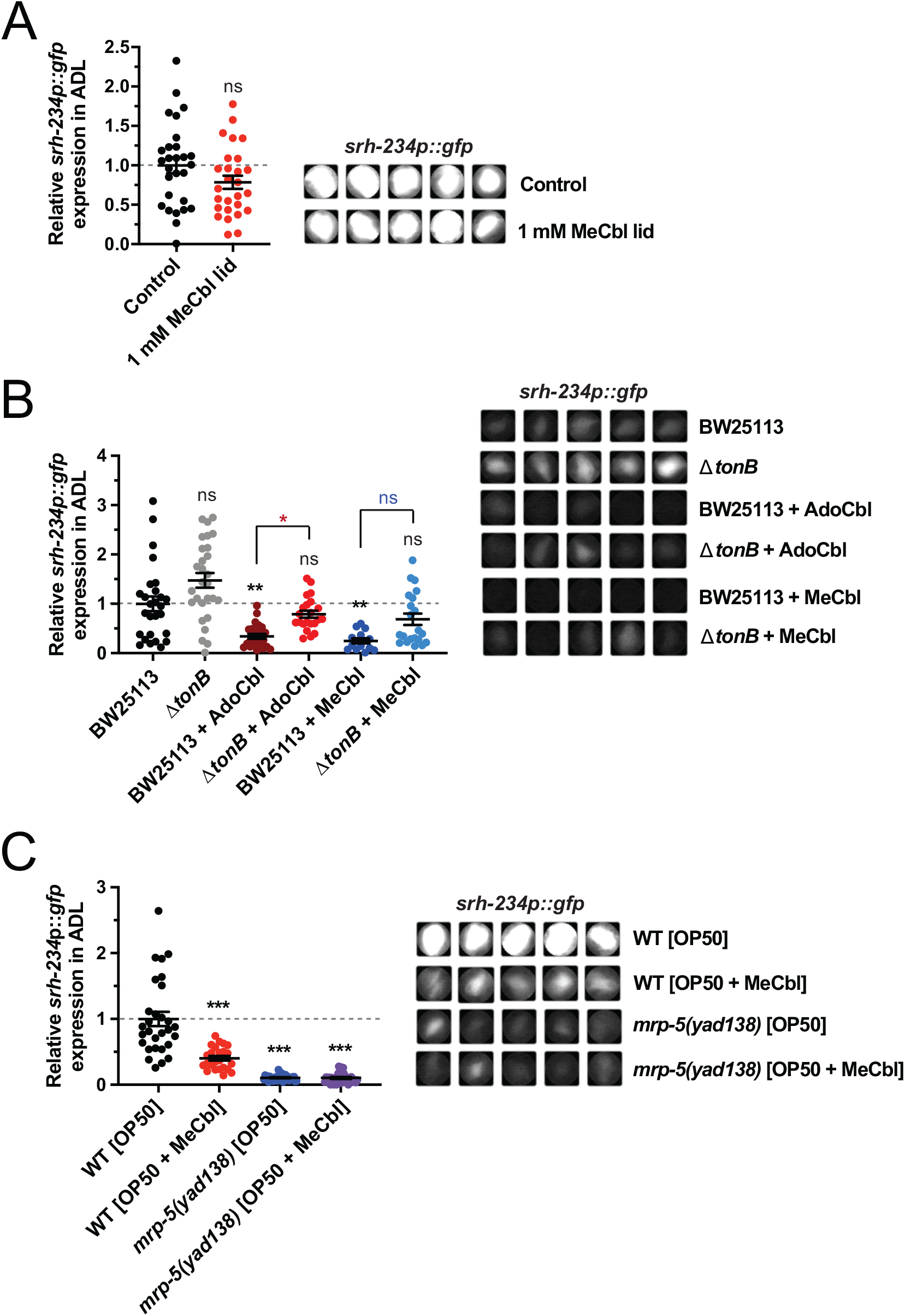
Bacterial vitamin B12 uptake regulates *srh-234* expression. **(A)** Relative expression of *srh-234p::gfp* in the ADL cell body of adult animals fed on OP50 diets on NGM agar plates covered with lids containing 1 mM MeCbl. Data are represented as the mean ± SEM (n>25 animals). ns, not significant by an unpaired 2-tailed *t*-test. **(B)** Relative expression of *srh-234p::gfp* in the ADL cell body of adults fed the *E. coli* Δ*tonB* mutant (strain JW5195) compared to its parental wild-type strain (BW25113) supplemented with or without AdoCbl and MeCbl (64 nm final concentrations). Data are represented as the mean ± SEM (n=14-28 animals for each condition). ** indicates values that are different from wild-type fed on *E. coli* OP50 or *Comamonas* DA1877 at *p*<0.01 by a Kruskal-Wallis test with Dunn multiple-comparisons test. Of note, *srh-234p::gfp* is weakly expressed on an *E. coli* BW25113 diet. **(C)** Relative expression of *srh-234p::gfp* in the ADL cell body of *mrp-5* mutants fed on *E. coli* OP50 diets supplemented with MeCbl (64 nM final concentration) compared to wild-type. Data are represented as the mean ± SEM (n>24 animals). *** indicates values that are different from wild-type fed on *E. coli* OP50 at *p*<0.001 by a Kruskal-Wallis test with Dunn multiple-comparisons test. **(A-C)** Right panels: Representative cropped images of *srh-234p::gfp* expression in the ADL cell body with the indicated compounds, genotypes and/or conditions. ns, not significant.

The *tonB* gene encodes a vitamin B12 transporter present in *E. coli* bacteria that allows these bacteria to import vitamin B12 from the extracellular environment (Bassford et al. 1976; Kadner 1990). When we exposed *C. elegans* to the *E. coli* K12-type BW25113 strain with loss-of-function mutations in *tonB*, animals showed a slightly increased *srh-234p::gfp* expression in ADL neurons compared to animals fed on wild-type *E. coli* BW25113 (**Fig. 4B**). As with OP50 diets, animals fed on *E. coli* BW25113 supplemented with either 64 nM AdoCbl or MeCbl significantly reduced *srh-234p::gfp* expression, which could be suppressed, at least in part, by *tonB* mutations (**Fig. 4B**). These results suggest that *E. coli* bacteria may likely function as the vehicle for vitamin B12 via the *tonB* transporter to regulate *srh-234* expression. However, alternate *tonB*-independent routes may be required as well to regulate *srh-234*. Consistent with food ingestion being the main vehicle for vitamin B12, we found that aztreonam-treated *Comamonas* DA1877 that *C. elegans* cannot eat but still smell and touch, partially suppresses the vitamin B12-mediated reduction of *srh-234* expression (**Fig. S4C**).

We next tested the role of the MRP-5 vitamin B12 transporter in *srh-234* regulation, which has been proposed to export vitamin B12 from the intestine to other tissues to support embryonic development of *C. elegans* (Na et al. 2018). We found that *mrp-5* mutations did not suppress the vitamin B12-mediated reduction in *srh-234* expression, although the interpretation of this negative result is confounded by the observation that *mrp-5* mutations strongly reduce *srh-234* expression in ADL when animals were fed the *E. coli* OP50 diet without vitamin B12 (**Fig. 4C**). Dye-filling of ADL was normal in *mrp-5* mutants (100% of animals dye-fill, n>25), suggesting that reduced *srh-234* expression in *mrp-5* mutants is not due to an ADL morphology defect (**Fig. 4D**). Thus, *mrp-5* may have additional yet unknown roles in *srh-234* regulation on the OP50 diet.

In summary, these data suggest that rather than directly sensing vitamin B12 levels, it is more likely that dietary-supplied vitamin B12 ingested by *C. elegans* regulates *srh-234* expression levels in ADL neurons.

### MEF-2 is required for the vitamin B12-mediated reduction in *srh-234* expression

To further interrogate the mechanisms underlying the vitamin B12-dependent regulation of *srh-234* gene expression in ADL neurons, we examined candidate components and pathways. We previously reported that the MEF-2 transcription factor acts together with bHLH factors to regulate the starvation-dependent regulation of *srh-234* expression (Gruner et al. 2016). In this mechanism, MEF-2 acts cell-autonomously with bHLH factors HLH-2/HLH-3 in ADL neurons, while HLH-30 and MLX-3 bHLH factors function in the intestine to non-cell-autonomously regulate *srh-234* expression in ADL in response to starvation signals. We found that a mutation in *mef-2* but not in *hlh-30* can fully suppress the vitamin B12-dependent reduction in *srh-234* expression when animals were fed a *Comamonas* DA1877 diet, suggesting that MEF-2 is required for the vitamin B12-dependent regulation of *srh-234* (**Fig. 5A, S5A**).

**Fig. 5:**
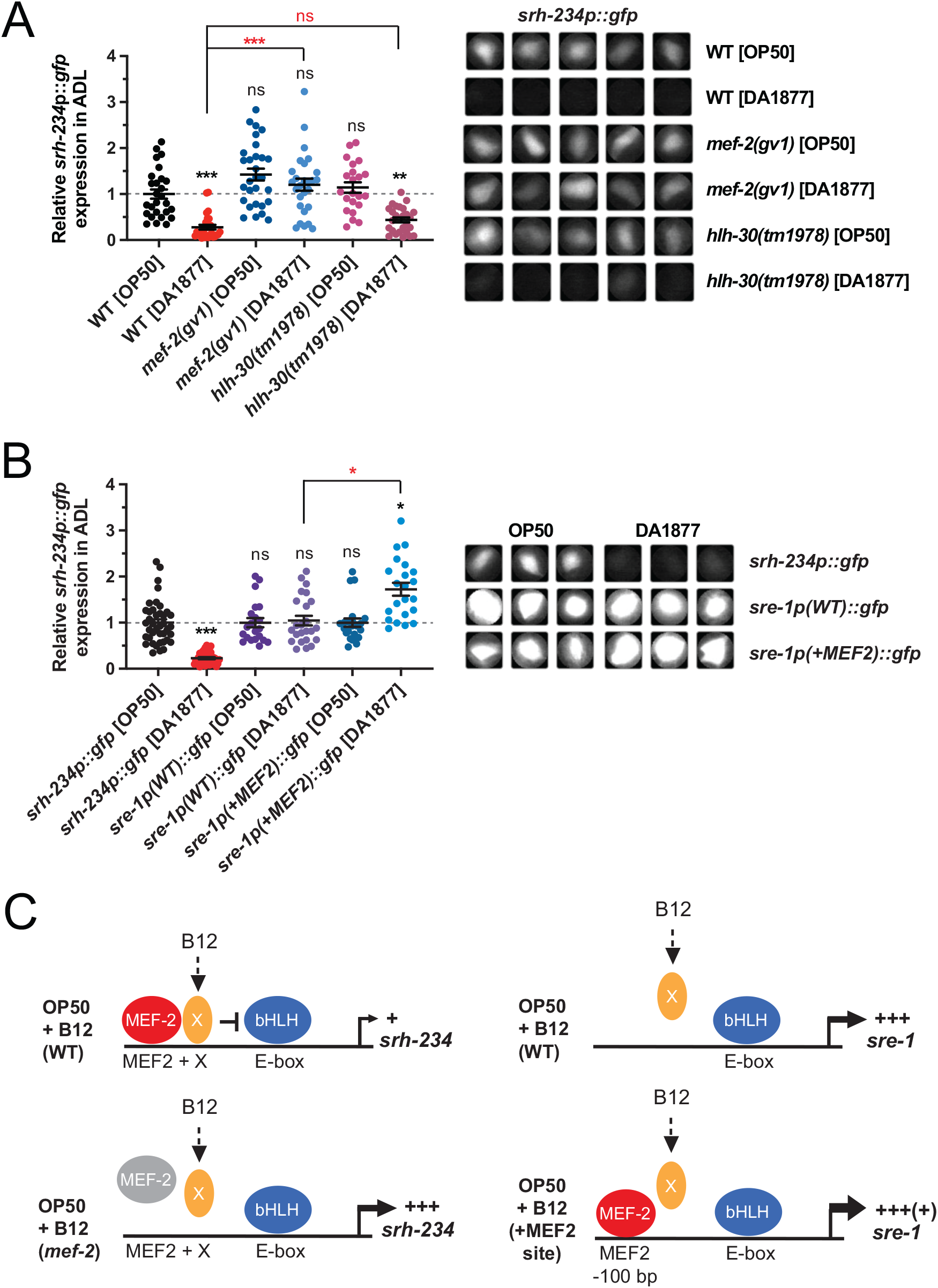
*mef-2* is required for vitamin B12-dependent regulation of *srh-234*. **(A)** Relative expression of *srh-234p::gfp* in the ADL cell body of *mef-2* and *hlh-30* mutants when adults are fed on *Comamonas* DA1877 compared to *E. coli* OP50 diets. Data are represented as the mean ± SEM (n>22 animals). ns, not significant. ***, ** indicates values that are different from wild-type fed on *E. coli* OP50 or *Comamonas* DA1877 at *p*<0.001 and *p*<0.01, respectively, by a Kruskal-Wallis test with Dunn multiple-comparisons test. Right panel: Representative cropped images of *srh-234p::gfp* expression in the ADL cell body with the indicated genotypes and diet conditions. Images were acquired at the same but lower exposure time. **(B)** Relative expression of wild-type *sre-1p::gfp* (*sre-1p(WT)::gfp*) or the *sre-1* promoter with the inserted MEF2 binding site sequence (*sre-1p(+MEF2)::gfp*) in the ADL cell body of adults fed on *Comamonas* DA1877 compared to an *E. coli* OP50 diet. Data are represented as the mean ± SEM (n=21-40 animals). ns, not significant. ***, * indicates values that are different from wild-type fed on *E. coli* OP50 or *Comamonas* DA1877 at *p*<0.001 and *p*<0.05, respectively, by a Kruskal-Wallis test with Dunn multiple-comparisons test. For *sre-1* expression, data were normalized to the *sre-1p::gfp* reporter fed on *E. coli* OP50. For *srh-234* expression, data were normalized to *srh-234p::gfp* fed on *E. coli* OP50. Right panel: Representative cropped images of *srh-234* or *sre-1* expression in the ADL cell body with the indicated diet conditions. **(C)** Model based on findings shown in panel B explaining the observed expression changes for *srh-234* in *mef-2* mutants, and *sre-1* with a *srh-234* MEF2-binding site inserted in its promoter upstream and close to the identified E-box that drives *sre-1* expression in the ADL neuron. +, +++, and +++(+) indicates low, high, and highly increased expression levels, respectively.

Since the *srh-234 cis*-regulatory region contains a MEF-2 binding site (**Fig. S5B**), which is required to repress but not to promote *srh-234* expression in starved conditions (Gruner et al. 2014), we examined whether this MEF-2 binding site was sufficient for the vitamin B12-dependent regulation of *srh-234* expression. To test this, we used a transgenic reporter strain of the *sre-1* promoter fused to *gfp* with or without the MEF-2 binding site identified in the *srh-234* promoter. The *sre-1* promoter is specifically and highly expressed in ADL neurons, but levels of *sre-1* expression are not changed by vitamin B12 (**Fig. S1D, 1E**). Surprisingly, we found that animals carrying a transgene of the *sre-1* promoter with the inserted MEF-2 site (*sre-1p(+MEF2)::gfp*) showed similar *sre-1* expression levels in ADL when fed *Comamonas* DA1877 compared to wild-type *sre-1p::gfp* animals (*sre-1p(WT)::gfp*) on the same diet (**Fig. 5B**). These results suggest that in contrast to the starvation-dependent regulation of *srh-234* (Gruner et al. 2014), insertion of the MEF2 binding site alone is not sufficient for the vitamin B12-dependent regulation of *srh-234* expression levels in ADL neurons, suggesting the requirement of another yet unknown factor that may act together with MEF-2 (**Fig. 5C**). Together, these findings show that the function of the MEF-2 transcription factor is necessary for regulation of *srh-234* mediated by dietary-supplied vitamin B12.

## DISCUSSION

In this study, we show that the expression levels of the *srh-234* chemoreceptor gene in the ADL sensory neuron type is regulated by dietary vitamin B12. In a low vitamin B12 *E. coli* diet, *srh-234* is highly expressed in ADL but not when *C. elegans* is fed a high vitamin B12-producing *Comamonas* diet (**Fig. 6**). This vitamin B12-mediated regulation of *srh-234* expression levels is dependent on the MEF-2 transcription factor. The mechanisms by which dietary vitamin B12 transcriptionally tunes *srh-234* could provide *C. elegans* the means to modify long-term changes in ADL-mediated responses.

**Fig. 6:**
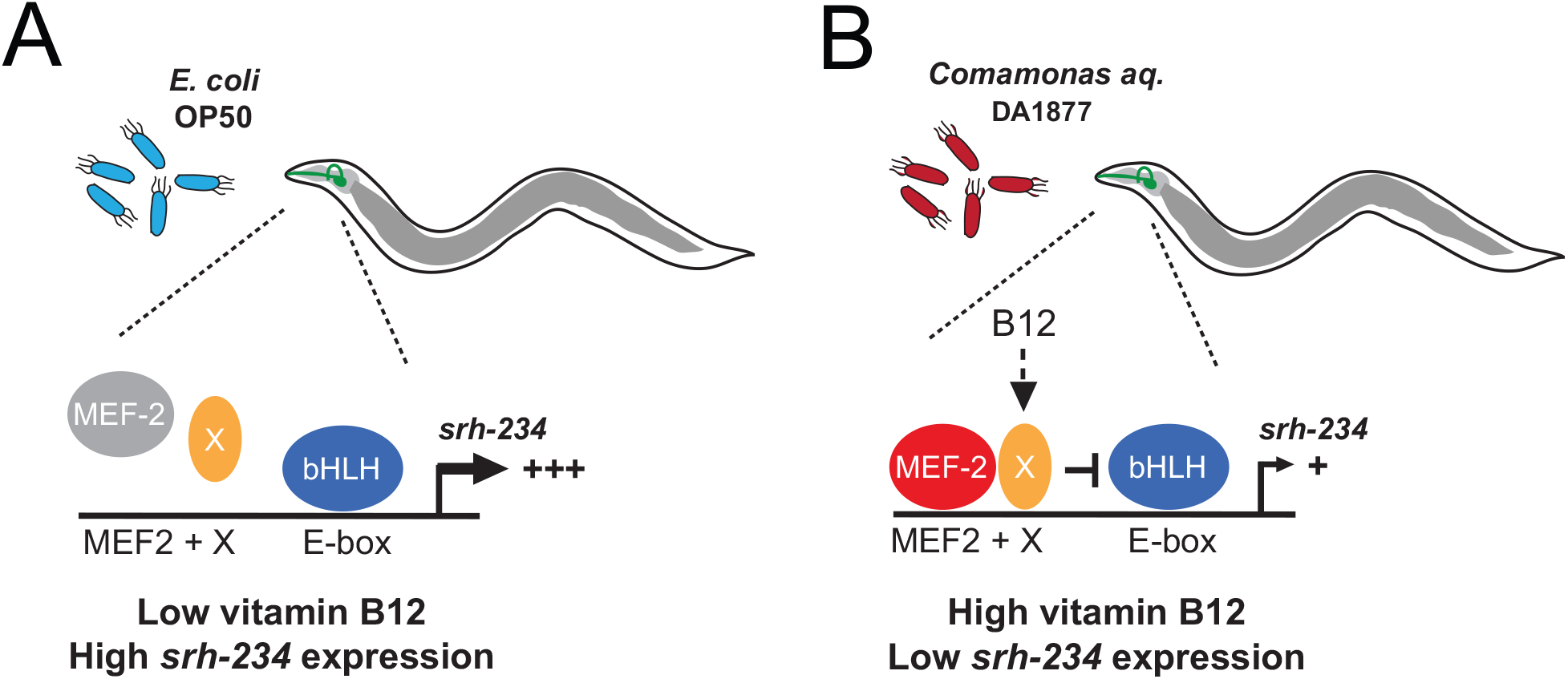
Model for the regulation of *srh-234* chemoreceptor expression levels in the ADL sensory neuron under different dietary conditions. Expression levels of *srh-234* in *C. elegans* animals is high (+++) when fed a low vitamin B12 diet of *E. coli* OP50 bacteria **(A)** but low (+) when fed a high vitamin B12 diet of *Comamonas aq*. DA1877 bacteria **(B)**. An unknown factor (X) may act together with the MEF-2 transcription factor to repress *srh-234* expression levels under conditions of high vitamin B12 via a bHLH/E-box module important to promote expression of *srh-234* in ADL.

This study complements our previous work (Gruner et al. 2014; Gruner et al. 2016), which explored the dynamics in *srh-234* expression upon starvation, which was dependent on MEF-2 function and its respective MEF2 binding site present in the *cis*-regulatory sequence of *srh-234*. Similarly, we show that loss-of-function *mef-2* mutations can suppress *srh-234* expression in ADL in response to feeding the vitamin B12-producing *Comamonas* bacteria, suggesting that MEF-2 has dual roles in regulating *srh-234* expression in response to both starvation and dietary vitamin B12. However, unlike starvation (Gruner et al. 2016), artificial introduction of the *srh-234* MEF2 binding site into the *cis*-regulatory sequence of the *sre-1* gene close to its ADL E-box site (McCarroll et al. 2005) did not confer vitamin B12-induced downregulation via MEF-2. Based on these findings, we propose a model (**Fig. 6**) in which animals fed a high vitamin B12 *Comamonas* diet reduce *srh-234* expression via a transcriptional module consisting of a MEF-2 factor and its respective MEF2 binding site, together with a yet unknown factor (X) stimulated by dietary vitamin B12. This in turn may repress bHLH factors through an E-box site that promotes *srh-234* in ADL neurons via a complex mechanism involving a combination of different bHLH heterodimer pairs (Gruner et al. 2016). When animals are fed a low vitamin B12 diet of *E. coli* OP50, MEF-2 activity no longer represses *srh-234* expression in ADL. Thus, MEF-2 activity is necessary for proper *srh-234* regulation in response to dietary-supplied vitamin B12, but the exact pathways by which vitamin B12 modulates *srh-234* remains to be discovered.

Our results also show that supplementing exogenous propionate to a *E. coli* OP50 diet, and mutations in the canonical (*pcca-1, pccb-1*) and shunt propionate (*acdh-1, hphd-1*) breakdown pathways, are able to override the repressing effects of vitamin B12 on *srh-234* expression. This may suggest that a toxic build-up of propionate levels in *C. elegans* regulates *srh-234* expression in ADL neurons, and that the balance between vitamin B12 and propionate levels is important for tuning the promoter activity of *srh-234*. In mammalian models of propionic acidemia, animals lacking the propionyl CoA-carboxylase (PCCA) were found to have elevated propionate levels shortly after birth (Miyazaki et al. 2001). Similarly, *pcca-1* mutant animals in *C. elegans* may have naturally elevated propionate levels that cannot be restored to normal levels by vitamin B12 sufficient diets alone (Watson et al. 2016). Consistent with a persistent accumulation of propionate in *C. elegans* modulating *srh-234* promoter activity, we show that *pcca-1* and *pccb-1* mutants significantly enhance the levels of *srh-234* expression on a *E. coli* OP50 diet that is unable to efficiently breakdown propionate by the canonical pathway. Conversely, *srh-234* expression levels are strongly reduced in ADL when exposed to low propionate levels; for instance, in animals that are food deprived (starved) or exposed to high vitamin B12 conditions. Studies in rats demonstrated that after two days of starvation, propionate levels are rapidly decreased but again restored after re-feeding (Illman et al. 1986).

The nociceptive ADL neuron where *srh-234* is specifically expressed mediates avoidance responses to a wide variety of environmental signals such as odors (Chao et al. 2004; Troemel et al. 1995; Troemel et al. 1997), pheromones (Jang et al. 2012), and heavy metals (Sambongi et al. 1999; Wen et al. 2020). Since chemoreceptor genes expressed in a specific chemosensory neuron type are generally linked to a common chemical response determined by the identity of the neuron in *C. elegans*, with a few exceptions in which neurons switch their preference towards odors (Tsunozaki et al. 2008), it is probable that the *srh-234* chemoreceptor may detect aversive chemical stimuli perceived by ADL. Interestingly, vitamin B12 in mammals has anti-nociceptive properties (Erfanparast et al. 2014), and the activity of certain olfactory receptors in tissues other than neurons can respond to propionate (Pluznick et al. 2013), which is a metabolic byproduct produced by gut bacteria in mammals (Morrison and Preston 2016). It is therefore tempting to speculate that vitamin B12 obtained through ingestion alters ADL-mediated nociceptive responses by changing the expression of individual chemoreceptor genes such as *srh-234*. However, nothing is known about whether vitamin B12 or propionate levels affects ADL-mediated responses in *C. elegans*. Other than growth, development, and lifespan (Bito et al. 2013; MacNeil et al. 2013), only recently vitamin B12 in the diet has been shown to be an important micronutrient in the regulation of predatory behaviors between nematodes (Akduman et al. 2020). In support of a model by which dietary vitamin B12 absorbed by the *C. elegans* intestine regulates *srh-234*, our previous work (Gruner et al. 2016) demonstrated that an intestine-to-ADL interaction is necessary to regulate *srh-234* expression as a function of feeding state. This communication between the intestine and the ADL neuron involves the action of non-cell-autonomous pathways including insulin-like peptides and *hlh-30*/*mxl-3* bHLH factors. We show that vitamin B12-mediated regulation of *srh-234* is not dependent on *hlh-30* function, suggesting that further research is needed to investigate how dietary vitamin B12 regulates *srh-234* expression in ADL neurons.

The *srh-234* chemoreceptor gene is one of a large repertoire of over 1,300 chemoreceptor genes (Robertson 2000), many of which are localized in a relatively small subset of chemosensory neurons (Vidal et al. 2018), such that each neuron expresses multiple chemoreceptor genes. We show here dynamic changes in the expression levels of the *srh-234* chemoreceptor gene localized in the ADL sensory neuron type in response to changing dietary vitamin B12. Other studies have illustrated that dynamic expression changes in individual chemoreceptor genes can have profound effects on behavioral outcomes. For instance, changes in the expression levels of the *odr-10* olfactory receptor required to sense diacetyl (Sengupta et al. 1996) in the male *C. elegans* contributes to its plasticity in food detection and feeding/exploration decisions in order to locate mates (Ryan et al. 2014). Further research will determine what the functional consequences are of the plasticity in *srh-234* chemoreceptor gene expression in ADL neurons in response to dietary vitamin B12.

## DATA AVAILABILITY

All data are available as part of this manuscript and are posted on the Open Science Framework (will be deposited). Supplemental data and information are available online.

## ACKNOWLEDGEMENTS

We thank Marian Walhout, the Keio Knockout collection, and the *E. coli* Genetic Stock Center for bacterial strains. We thank Dong Yan for the *mrp-5* mutant strain. We thank the *Caenorhabditis elegans* Center, which is funded by the National Institutes of Health Office of Research Infrastructure Programs (P40 OD010440), for both *C. elegans* and bacterial strains. We also thank the Cellular and Molecular Imaging Core facility of the COBRE Integrative Neuroscience Center at the University of Nevada for providing equipment and resources.

## FUNDING

This work was supported by the National Science Foundation grant IOS 1353014 to A.M.V., and the National Institutes of Health grant R01 NS107969 to A.M.V. Research reported in this work used the Cellular and Molecular Imaging (CMI) core facility supported by the National Institute of General Medical Sciences of the National Institutes of Health grant P20 GM103650.

## CONFLICTS OF INTEREST

None declared.

## Notes

### Competing Interest Statement

The authors have declared no competing interest.

