## Supplemental Data for "Dietary vitamin B12 regulates chemosensory receptor gene expression via the MEF2 transcription factor in *Caenorhabditis elegans*"

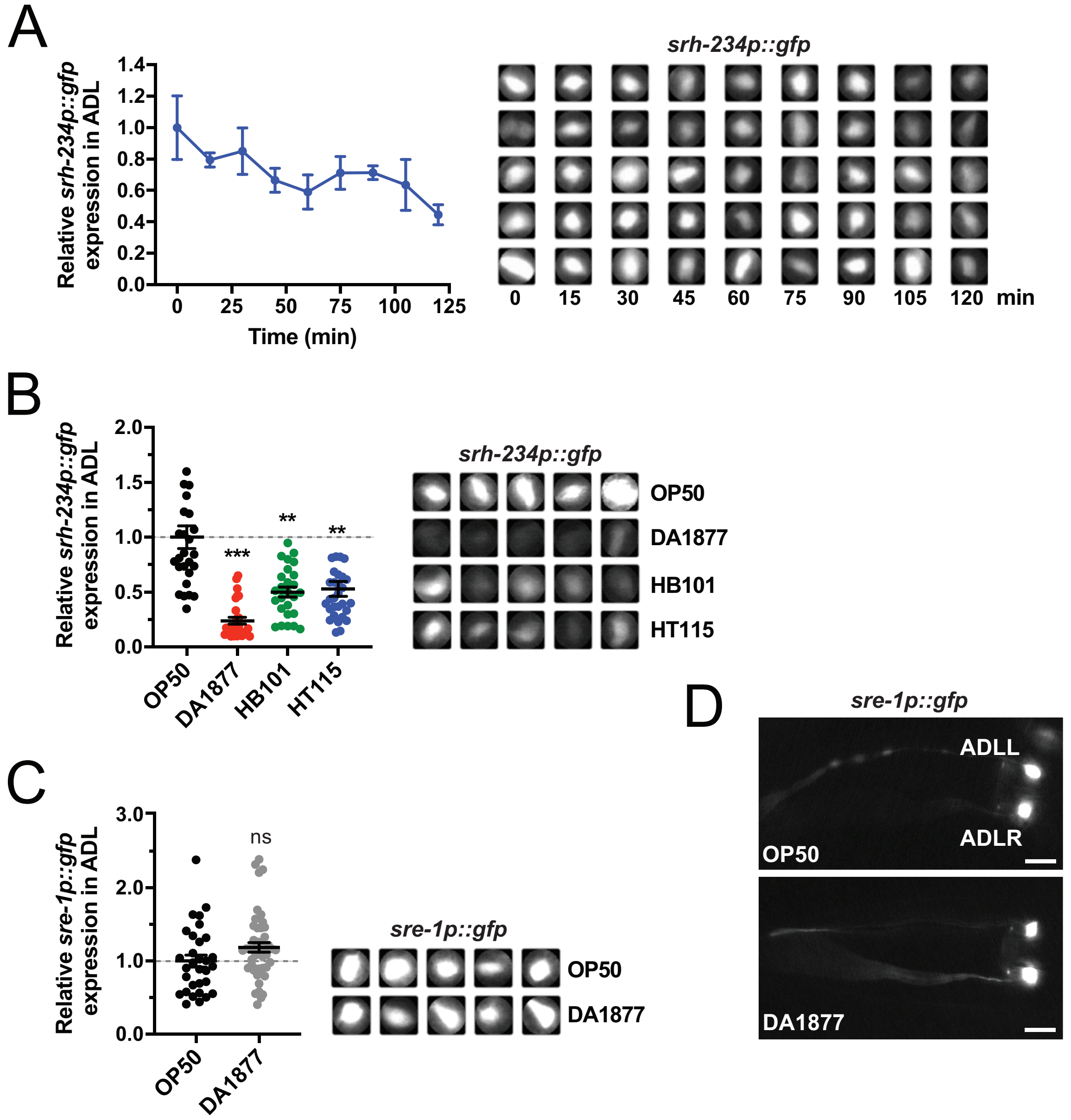


**Fig. S1: *srh-234* expression in ADL neurons is rapidly downregulated in the presence of *Comamonas* DA1877.**
**(A)** Time-course of the relative *srh-234p::gfp* expression levels in the ADL cell body of adult animals when transferred from an *E. coli* OP50 diet to a *Comamonas* DA1877 diet. Expression of *srh-234p::gfp* in adults was measured each 15 min at the same exposure time. The graph shows the mean ± SEM (n=5 animals for each timepoint). Right panel: Representative cropped images of *srh-234p::gfp* expression in the ADL cell body in adult animals fed DA1877 at the indicated exposure times. **(B)** Relative expression levels of *srh-234p::gfp* in the ADL cell body of adults fed *E. coli* HB101 and HT115 diets compared to *E. coli* OP50 and *Comamonas* DA1877. Data are represented as the mean ± SEM (n>25 animals for each diet). ***, ** indicates values that are different from wild-type fed on *E. coli* OP50 at $p<0.001$ and $p<0.01$, respectively, by a

Kruskal-Wallis test with Dunn multiple-comparisons test. **(C)** Relative expression of *sre-1p::gfp* in the ADL cell body of adult animals fed OP50 and DA1877 diets. Animals contain stably integrated copies of a *sre-1p::gfp* transgene (*otIs24*). Data are represented as the mean ± SEM (n>25 animals). ns, not significant by an unpaired 2-tailed *t*-test. **(D)** Representative images of *sre-1p::gfp* expression of *E. coli* OP50- and *Comamonas* DA1877-fed animals are a ventral view of ADL sensory neurons, and images were acquired at the same exposure time. Scale is 15 µm. **(A-D)** Images were acquired at the same exposure time.

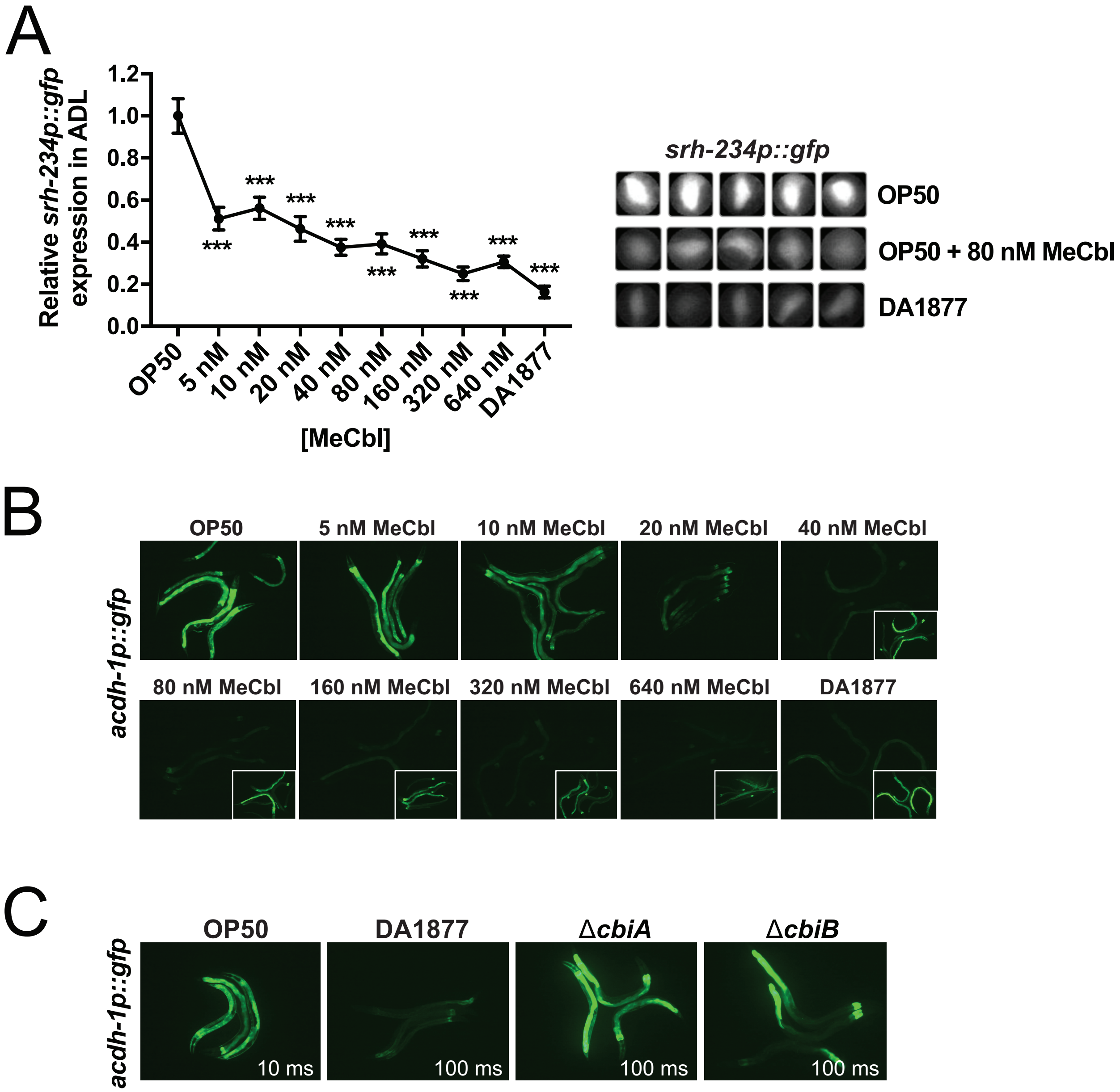


**Fig. S2: Dose dependent regulation of *srh-234* expression by vitamin B12.** **(A-B)** Dose-dependent decrease in *srh-234p::gfp* expression in the ADL cell body (A) and *acdh-1p::gfp* expression in the intestine **(B)** of *E. coli* OP50-fed animals supplemented with increasing concentrations of MeCbl. The graph shows the mean ± SEM (n>12 animals for each time-point). *** indicates values that are different from wild-type animals fed on *E. coli* OP50 at *p*<0.001 performed by a 2-way ANOVA with Tukey multiple-comparisons test. Right Panel: Representative cropped images of *srh-234p::gfp* expression in the ADL cell body. **(B-C)** The dietary sensor *acdh-1p::gfp* was used as a positive control for the action of different MeCbl concentrations (B) and when animals were fed Δ*cbiA* and Δ*cbiA* mutant bacteria of *Comamonas aq.* **(C)**. **(B-C)** Images were acquired at the same exposure time for comparison unless indicated otherwise, and insets in panel B are images taken at a higher exposure time.

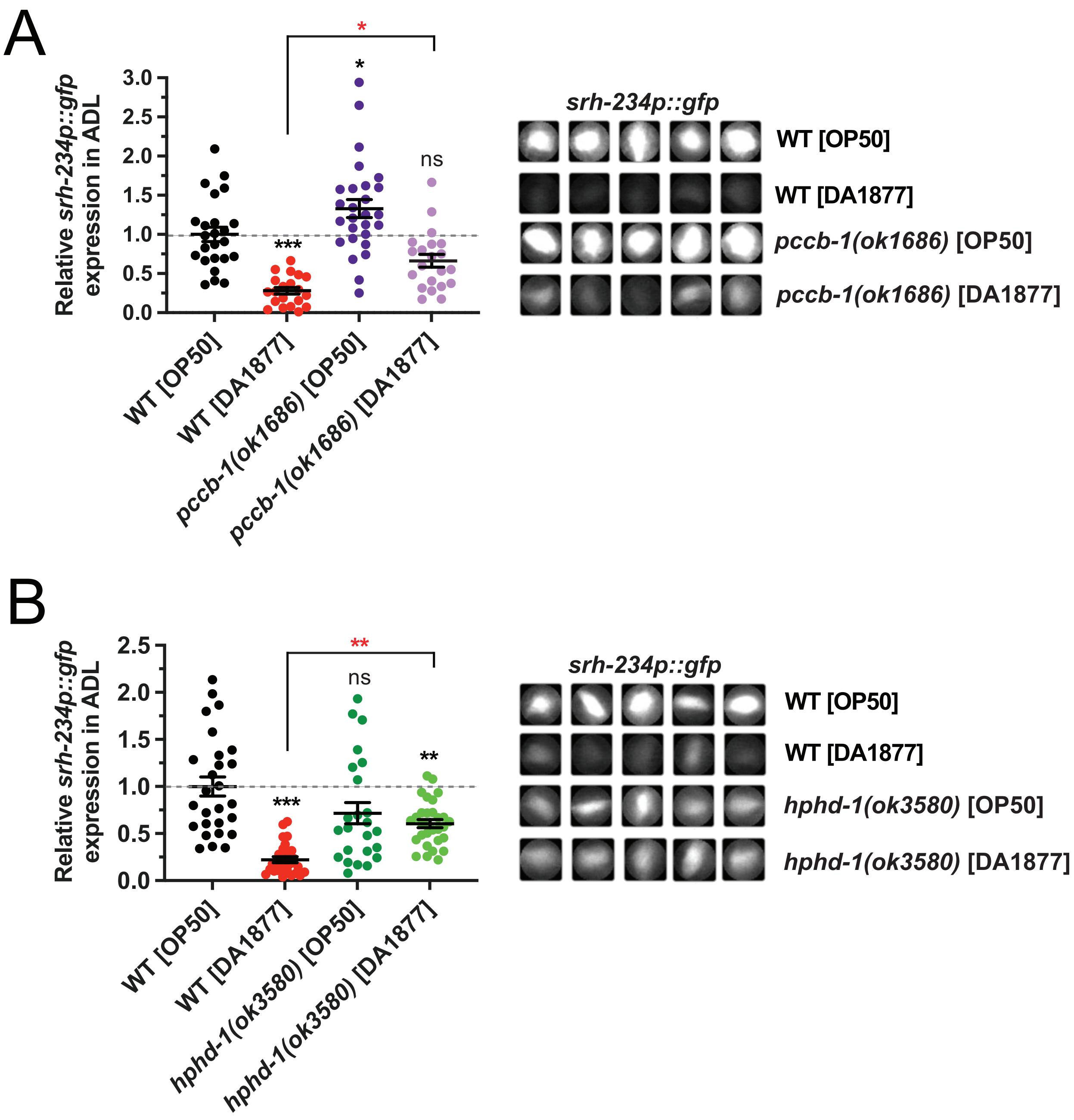


**Fig. S3: Vitamin B12 reduces *srh-234* expression but not in animals with mutations in the *pccb-1* and *hphd-1* genes.**
Relative expression of *srh-234p::gfp* in the ADL cell body of adult animals with mutations in the canonical propionate breakdown pathway gene *pccb-1* (propionyl-CoA carboxylase) **(A)** and the propionate shunt gene *hdpd-1* (3-hydroxypropionate-oxoacid transhydrogenase) **(B)** when fed either *E. coli* OP50 or *Comamonas* DA1877 diets. n>25 animals for each condition. **(A-B)** Data are represented as the mean $\pm$ SEM (n>20 animals). ns, not significant, ***, ** indicates values that are different from wild-type fed on *E. coli* OP50 or *Comamonas* DA1877 at $p<0.001$ and $p<0.01$, respectively, by a one-way ANOVA with Tukey multiple-comparisons test.

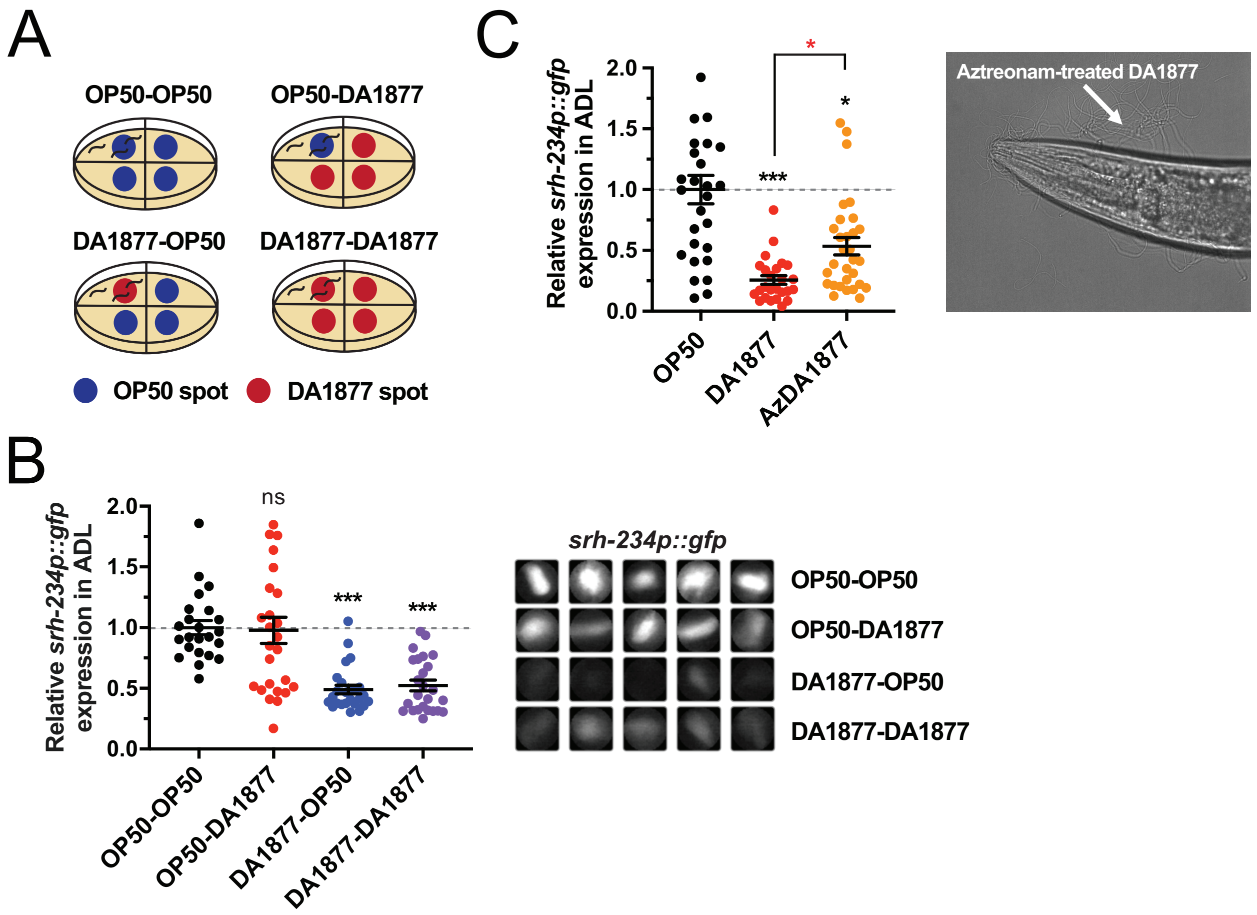


**Fig. S4: Sensory signals of vitamin B12 do not regulate *srh-234* expression.** **(A)** Schematic of a bacterial olfactory assay set-up using quadrant assay plates. Adult animals are placed into one quadrant allowing only one diet to be ingested, while surrounding quadrants are seeded with either the *E. coli* OP50 diet (control) or the *Comamonas* DA1877 diet. **(B)** Relative expression of *srh-234p::gfp* in the ADL cell body of adults fed on either the *E. coli* OP50 or *Comamonas* DA1877 diet on quadrant plates with surrounding inaccessible diets. Data are represented as the mean ± SEM (n>23 animals for each diet). ns, not significant. *** indicates values that are different from wild-type fed on *E. coli* OP50 (OP50-OP50) at $p<0.001$ by a Kruskal-Wallis test with Dunn multiple-comparisons test. **(C)** Relative expression of *srh-234p::gfp* in the ADL cell body of adults fed with *E. coli* OP50 (OP50), *Comamonas* DA1877 (DA1877) and aztreonam-treated *Comamonas* DA1877 (AzDA1877). Data are represented as the mean ± SEM (n>25 animals for each diet). ***, * indicates values that are different from wild-type fed on *E. coli* OP50 or *Comamonas* DA1877 $p<0.001$ and $p<0.01$, respectively by a Kruskal-Wallis test with Dunn multiple-comparisons test. Right image shows a *C. elegans* head surrounded by aztreonam-treated DA1877 bacteria that grow in long chains that *C. elegans* cannot eat due to its large size.

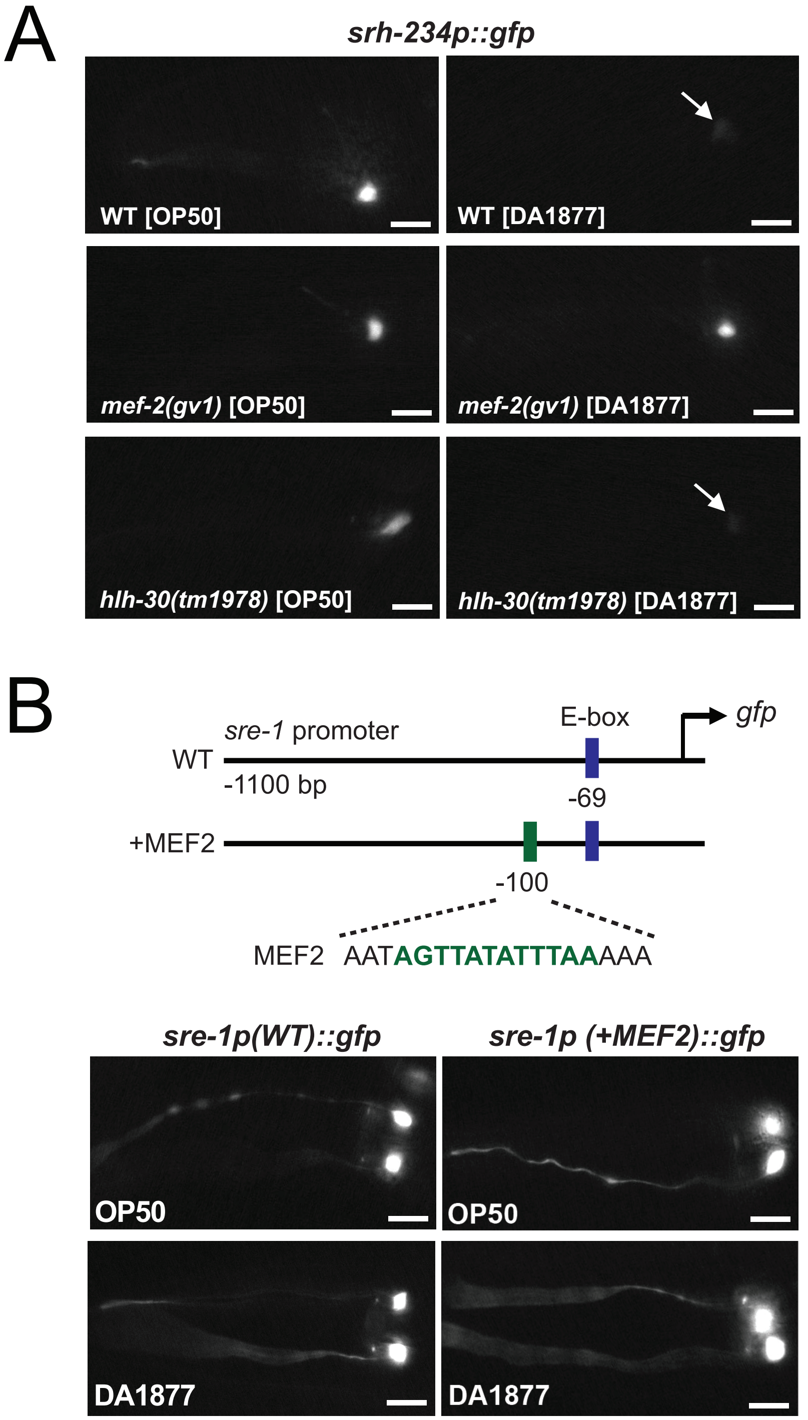


**Fig. S5: Introduction of the *srh-234* MEF2 site in the *cis*-regulatory region of *sre-1* does not confer regulation by vitamin B12.**
(A) Representative images of *srh-234p::gfp* expression in the ADL cell body of *mef-2* and *hlh-30* mutants fed on *Comamonas* DA1877 compared to *E. coli* OP50 diets. Images were taken at the same exposure time (anterior is left). Scale is 15 μm. (B) The indicated lengths and positions of predicted regulatory elements relative to the translational start site of *sre-1* fused to the *gfp* coding sequence in an expression vector.

Sequences in blue and green indicate the predicted E-box motif of *sre-1* with the inserted MEF2 binding site sequence of *srh-234*, respectively. Lower panel: Representative expression of ADL sensory neurons driven by wild-type *sre-1 cis*-regulatory sequences without (*sre-1p(WT)::gfp*) or with the MEF2 binding site of the *srh-234* promoter inserted (*sre-1p(+MEF2)::gfp*) in adult wild-type animals fed on *Comamonas* DA1877 compared to *E. coli* OP50. Images are ventral views of ADL sensory neurons with left and right cell bodies of ADL (anterior is left), and were acquired at the same exposure time. Scale is 15 µm.
